# Proteostasis governs differential temperature sensitivity across embryonic cell types

**DOI:** 10.1101/2022.08.04.502669

**Authors:** Michael W. Dorrity, Lauren M. Saunders, Madeleine Duran, Sanjay R. Srivatsan, Brent Ewing, Christine Queitsch, Jay Shendure, David W. Raible, David Kimelman, Cole Trapnell

**Affiliations:** Department of Genome Sciences, University of Washington, Seattle, WA 98195; Structural and Computational Biology, European Molecular Biology Laboratory, Heidelberg, DE 69117; Department of Biochemistry, University of Washington, Seattle, WA 98195; Department of Biological Structure, University of Washington, Seattle, WA 98195; Brotman Baty Institute for Precision Medicine, Seattle, WA 98195; Howard Hughes Medical Institute, Seattle, WA 98195

**Keywords:** developmental robustness, variability, zebrafish, single cell RNA-seq

## Abstract

The genetic program of embryonic development is remarkably robust, but temperature stress can degrade its ability to generate animals with invariant anatomy. While the stereotyped, consistent phenotypes associated with environmental stress during vertebrate development suggest that some cell types are more sensitive to stress than others, the basis of this sensitivity is unknown. Here, we characterize hundreds of individual zebrafish embryos under temperature stress using whole-animal single cell RNA-seq to identify cell types and molecular programs within them that drive phenotypic variability. We find that temperature perturbs the normal proportions and gene expression programs of numerous cell types and also introduces asynchrony in their development. The notochord is particularly sensitive to temperature stress, which we show is due to a specialized cell type, sheath cells. Further analyses show that sheath cells accumulate misfolded protein at elevated temperature, leading to a cascading structural failure of the notochord and irreversible anatomic defects in the embryo. Our study demonstrates that whole-animal single cell RNA-seq can characterize mechanisms important for developmental robustness and pinpoint molecular programs within specific cell types that constitute key failure points.

## Introduction

Temperature and developmental rate are commonly correlated across animal development ^1–3^. The acceleration of development at higher temperatures has been attributed to increased metabolic rate and protein synthesis ^4^. Within a species-specific tolerated range, embryos raised in elevated temperatures are phenotypically normal, suggesting that the developmental program is synchronously accelerated. While the mechanisms underlying this synchrony are not fully understood, recent studies have emphasized the role of proteostasis—the maintenance of proper protein folding, synthesis, and degradation—in determining developmental rate differences across species ^4–6^. Mechanisms regulating cellular proteostasis are also key for understanding the relationship between environmental stress and phenotype ^7^. Regulation of protein folding and synthesis is sensitive to diverse environmental signals ^8^ and misregulation of these processes has phenotypic consequences ^9–12^. Because the burden of proteostasis varies considerably across cell types ^13,14^, we sought to use transcriptional signatures of proteostasis to investigate how each lineage is able to buffer the effects of temperature stress. Perturbation of proteostasis during early development introduces phenotypic variability ^12,15,16^, but these phenotypes are non-random, suggesting that some cell types or developmental processes may be more sensitive to stress than others.

Here we investigate developmental robustness using zebrafish embryos, which are naturally exposed to temperature fluctuation, and display temperature-induced phenotypes and alterations in developmental rate. Specifically, we sought to understand whether cell types respond differently to temperature stress. Single cell genomic techniques can resolve heterogeneity in developmental timing and in levels of molecular processes promoting proteostasis across cell types, but these techniques are limited in their ability to capture variability in the form of biological replicates due to the standard practice of pooling embryos. To overcome this, we use single-cell combinatorial indexing combined with DNA oligo hashing to capture transcriptional states and cell type abundances for hundreds of individual zebrafish embryos at multiple temperatures and time points. We leverage this large number of replicates to make statistical inferences about the source of environmentally-induced phenotypes, as well as to identify molecular sources and cell type-specific contributions to the loss of developmental robustness. From these analyses, we find that temperature affects both developmental rate and synchrony among cell types. Furthermore, temperature stress alters cell type composition, leaving a lasting imprint on the embryo that cannot be explained by differences in developmental stage. Finally, we show that perturbed proteostasis reveals cell type-specific temperature sensitivity in the notochord. Taken together, we identify cell type-specific mechanisms of developmental robustness by integrating individual-to-individual variation and accounting for differential responses to temperature across cell types.

## Results

### Multiplexed single-cell RNA-seq profiles individual embryos developing under temperature stress

To measure effects of temperature on developmental robustness, we raised zebrafish embryos over a range of increasingly stressful temperatures. In addition to the standard temperature of 28°C, we raised embryos in the standard condition as well as at two elevated temperatures, 32°C and 34°C. At elevated temperatures, embryos develop at a faster rate ^17^, with a substantial fraction of individuals showing axial defects of varying severity, along with other previously documented phenotypes (**Fig. 1a**) ^1,18–20^. Because we can profile individuals, we included embryos raised at elevated temperatures that were either phenotypically normal or had severe, temperature-induced phenotypes. To identify individual embryos, we used nuclear “hashing” wherein polyadenylated DNA oligos with unique barcodes for each embryo are added and fixed along with liberated nuclei after whole-embryo dissociation (**Fig. 1b**) ^21^. Treated and control embryos from three time points (24, 30, 36 hours post-fertilization [hpf]) were dissociated and hashed individually and subsequently processed with a single-cell combinatorial indexing protocol to isolate single nuclei with quality comparable to previous sci-RNA-seq experiments (**Extended Data Fig. 1b**) for transcriptome profiling ^22^. We recovered 5-10% of the cells (**Extended Data Fig. 1a-c**) from each individual embryo profiled (assuming 20,000-40,000 cells per embryo), which is sufficient to capture all major cell types for each individual. By combining the data from all 288 profiled individuals (403,992 total cells) we built a comprehensive atlas of cell type-specific responses to temperature stress (**Fig. 1c, Extended Data Fig. 1c**). We projected our temperature perturbation data onto our new annotated zebrafish development reference dataset produced from analyzing 1.2 million cells from 18 hpf to 96 hpf (**see accompanying manuscript, Saunders et al. 2022**) and identified 85 distinct cell types at these stages. Consistent with the robust progression of development despite elevated temperature, all expected cell types were represented in embryos raised at 32°C and 34°C, and we observed no global activation of the heat shock response (**Extended Data Fig. 1d**), which can diminish cell type-specific expression patterns ^23^.

**Figure 1.**
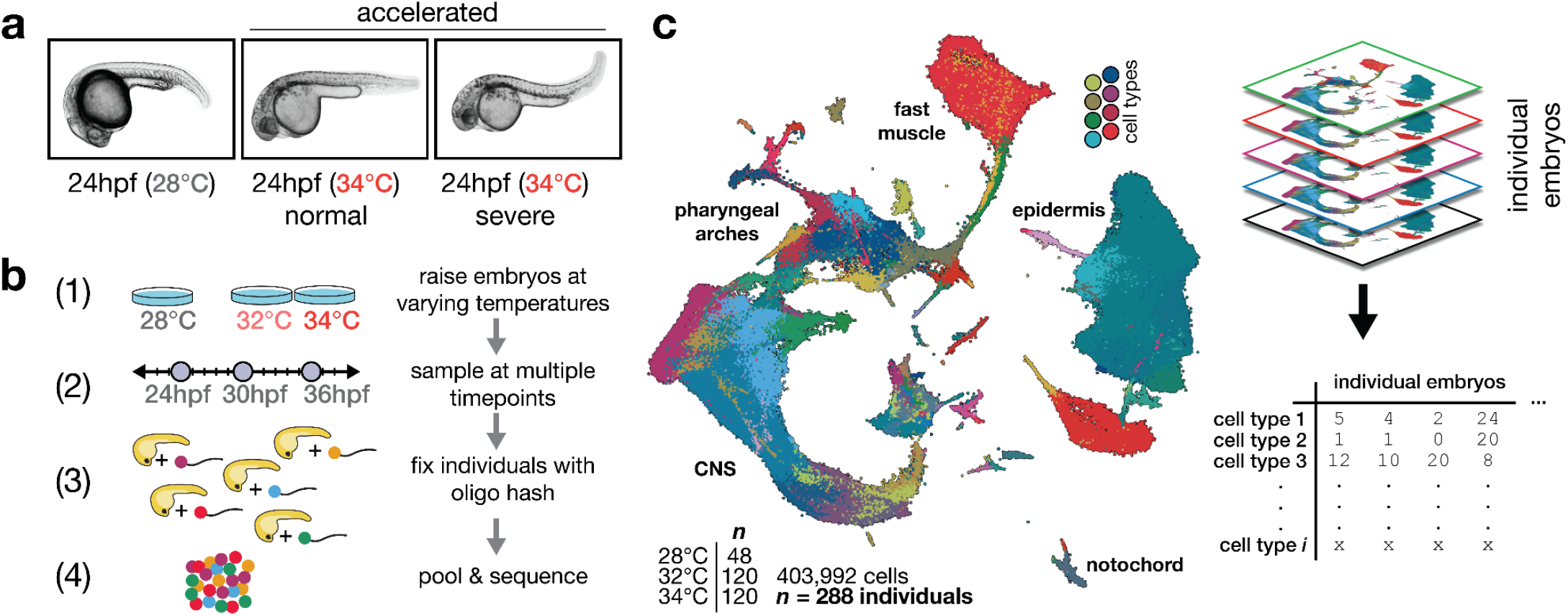
Effects of stress on phenotypic variability is captured via individual animal hashing of single-cell transcriptomes. **a**, Representative images of 24 hpf embryos raised at standard and elevated temperature; individual embryos with normal-looking and bent-tail phenotypes were included in the dataset. **b**, Experimental workflow for temperature perturbation experiment and individual embryo hashing. **c**, UMAP of temperature-perturbation dataset, projected into coordinate space of reference atlas (**see accompanying manuscript, Saunders et al. 2022**). Right side shows how single cell data are transformed to generate cell composition matrices.

### Variation in developmental stage can be determined from cell type composition

Because of variation in developmental progression over time and temperature, zebrafish studies categorize embryos based on a “staging series” of visible landmarks evident during development at standard temperature. For example, the proper stage-matched control for an embryo raised to 24 hpf at 34°C would be a 31 hpf embryo at 28°C ^17^. We therefore sought to develop a method to quantify the degree to which an embryo is accelerated directly from our single-cell data, as staging each embryo is essential for isolating the effects of temperature on cell type composition.

Using our reference dataset (**see accompanying manuscript, Saunders et al. 2022**), we analyzed individual embryos in two dimensions; embryos were grouped according to similarity in cell type composition, and this grouping produced a trajectory defined primarily by sample time point (**Fig. 2a**, middle, landmarks labeled on left). Individual temperature-treated embryos were projected into this low dimensional space and assigned a “pseudostage” value based on their relative position in the trajectory. Because temperature treatment distorts the relationship between clock time and embryo stage, we next tested whether pseudostaging captured the expected increases in developmental rate at high temperature. We projected high temperature embryos onto the reference embryo UMAP and found that treated embryos progressed further along the pseudostage trajectory than contemporaneous controls (**Fig. 2a, right**). Pseudostage values were strongly correlated (R^2 = 0.82) to acceleration predicted by a proposed linear model of temperature and developmental stage (**Fig. 2b**, see Methods) ^17^. Pseudostage values were also strongly correlated with whole-embryo stage values computed from nearest-neighbor average time points in individuals, rather than whole embryo cell type composition (**Fig. 2c**). Thus, whole-embryo cell type composition data captures variation in developmental stage, including that found in untreated embryos sampled at the same clock time (**Fig. 2b**). Staging embryos based on cell type composition is therefore sufficient to control for both biological and technical variation in developmental progression.

**Figure 2.**
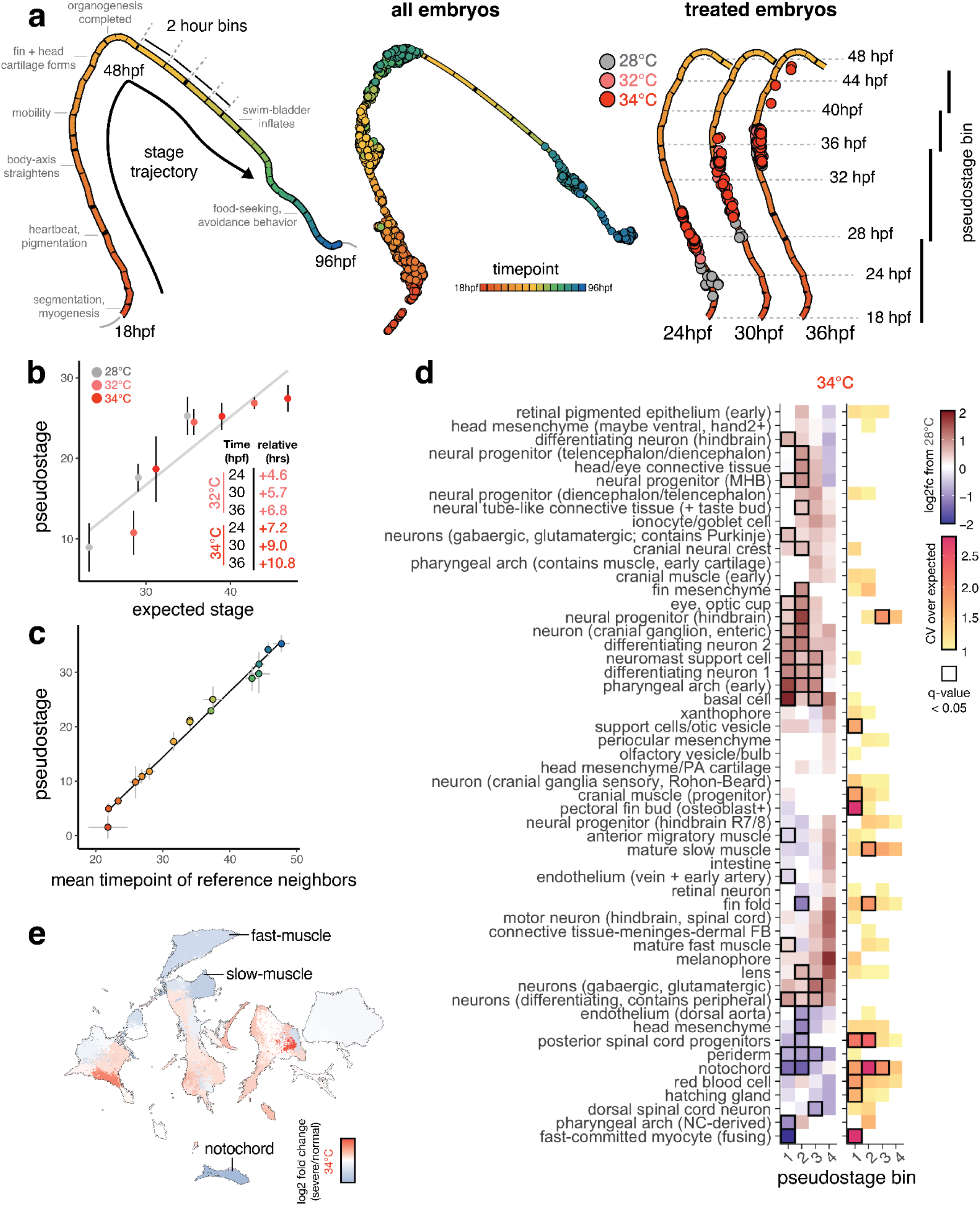
Staging embryos by cell type composition captures temperature-induced acceleration of development and allows isolation of temperature-dependent effects on cell abundance. **a**, Embryo trajectory produced using all individual embryos from the developmental reference; each segment of the principal graph represents a 2-hour window of development, with key events noted. Right panel shows how, at each time point, embryos from elevated temperature are “ahead” of their 28C counterparts on the embryo stage trajectory. **b**, Scatterplot showing mean pseudostage values are correlated with expectations from a linear model of temperature-induced acceleration of developmental rate. **c**, Scatterplot showing mean pseudostage values for all embryos in the reference dataset compared to a nearest-neighbor label transfer in transcriptome space; both cell composition and transcriptomes contain ample information on developmental stage. **d**, Heatmap showing the effects of temperature on mean cell type abundance relative to untreated, stage-matched controls (left) and on variability (CV relative to controls) of cell counts (right). Significant tests (q < 0.05) from beta binomial regression are indicated with a black box. Each column represents a pseudostage bin, wherein embryos from untreated and treated samples are stage-matched. **e**, UMAP projection of all cell types, colored by relative abundance change in severe (bent) individuals raised at 34°C compared to normal-looking embryos also raised at 34°C.

### Temperature stress alters cell type proportions and injects variability into embryogenesis

Next, we sought to explicitly evaluate the assumption that cell types develop synchronously and in proper proportions in embryos raised at different temperatures. Specifically, we wondered whether we could detect cell type-specific sensitivity to temperature in cell abundance data, as this may contribute to the stereotyped phenotypes that arise in embryos raised at high temperature. We grouped control and treated embryos of similar developmental stage into four pseudostage bins (**Fig. 2a, right**). Using both absolute cell counts, and relative proportions of each cell type per embryo, we performed a regression test using the beta binomial distribution, which is well-suited for compositional data ^24,25^. After accounting for differences in embryo stage in the model due to the different temperatures, we identified 20-30 cell types that showed significant (q-value < 0.05) increases or decreases in abundance with temperature (**Fig. 2d, left, Supplementary Table 1**), with a greater number of affected cell types at 34°C than 32°C. (**Extended Data Fig. 2a**). Of these, the most strongly affected cell types were notochord, dorsal aorta, and head mesenchyme, showing large reductions in cell number. Surprisingly, neural progenitor cell types showed significant increases in abundance. Other cell types such as periocular mesenchyme, xanthophore pigment cells, and red blood cells did not show significant changes in abundance as a function of temperature, suggesting that these cell types are not as sensitive to stress (**Fig. 2d)**. The effects of 32°C and 34°C treatments on cell abundances were correlated (R^2 = 0.69), suggesting that these sub-heat shock temperatures affected developmental processes similarly. Overall, many cell types showed consistent changes in proportion at higher temperatures, even after accounting for developmental stage, reinforcing the notion that stress fundamentally alters cell composition in embryos, rather than perturbing development of just a few lineages.

To test whether elevated temperature increased variability in cell composition globally, we compared multinomial models informed by pseudostage alone, or by pseudostage and temperature. We found 42% of the variation in whole-embryo cell composition between 24 and 36 hpf was explained by sample time point, with temperature explaining an additional 12% (**Extended Data Fig. 2c**), confirming a global increase in anatomical variability. To isolate specific sources of variability during embryogenesis, we identified cell types whose counts showed increased variance over a control expectation (shown as relative coefficient of variation [CV]). After correcting for the dependence of variance on the mean abundance of each cell type, we identified eleven cell types showing significant increases in variability at one of the elevated temperatures (**Fig. 2d, examples in Extended Data Fig. 2d, Supplementary Table 2**). However, within cell types, the largest dilations in CV occurred in a pseudostage-specific manner (**Fig. 2d, right**). For example, counts of posterior spinal cord progenitors showed increased variability at high temperature in earlier stages, but stabilized at later stages (**Fig. 2d, inset**). Across all cell types on average, relative CV values increased at higher temperatures (**Extended Data Fig. 2e**) and peaked between pseudostages 20-25, consistent with the previous observation that some developmental stages may be more sensitive to stress (**Extended Data Fig. 2e, upper panel**) ^26^.

We observed phenotypes of varying severity in embryos raised at high temperatures (severe phenotypes, **Fig. 1a**). We next sought to test whether altered cell composition contributed to differences in phenotypic severity. For embryos raised at 34°C, we classified the phenotype severity for each embryo (**Supplementary Table 3**), and, within a single temperature, compared cell type compositions of severe embryos to normal embryos. Embryos with more severe phenotypes showed reduced abundance of several cell types, including notochord, muscle, fin mesenchyme, pharyngeal arch (**Fig. 2e, Extended Data Fig. 2b**), suggesting that variation in cell type composition contributes to overall phenotypic variation, even within a temperature. Profiling cell type-specific sources of phenotype severity highlights the value of profiling large numbers of replicate embryos. Sensitive detection of temperature-dependent effects on cell abundances would not be possible with just a few replicate embryos, as resultant phenotypes can be obscured by individual-level phenotypic variability.

In summary, we find that some, but not all, cell types show altered abundance as a function of temperature, suggesting the response to temperature is integrated non-uniformly across cell types. This unexpected result raises the question: what is the molecular basis for these cell type-specific effects?

### Temperature introduces asynchrony in developmental rate across cell types

We next explored whether the temperature-dependent changes in cell type composition were attributable to changes in cell type-specific developmental rates. While the embryo as a whole develops 4-6 hours more quickly when raised at 32°C/34°C than at the standard 28°C (**Fig. 2b, inset**), variation in this acceleration across cell types has not previously been measured. To determine whether cell type-specific developmental rates vary, we examined relative differences in each cell type’s transcriptional “age,” a measure derived from comparison to our developmental reference, in temperature treatments compared to controls (**Extended Data Fig. 3a**, see Methods). For example, notochord cells in embryos raised at 32°C and 34°C showed +1.6hr and +2.2hr acceleration of transcriptional age relative to the whole-embryo expectation (**Extended Data Fig. 3b**). Several additional cell types, including the dorsal aorta, melanophore and hypochord cells (**Fig. 3a**, middle), showed a similar relative advancement, and we classified these cells as highly accelerated (**Fig. 3b**). In contrast, basal cells, pigment progenitors, and fin fold cells were unresponsive or decelerated at high temperature (**Extended Data Fig. 3b**).

**Figure 3.**
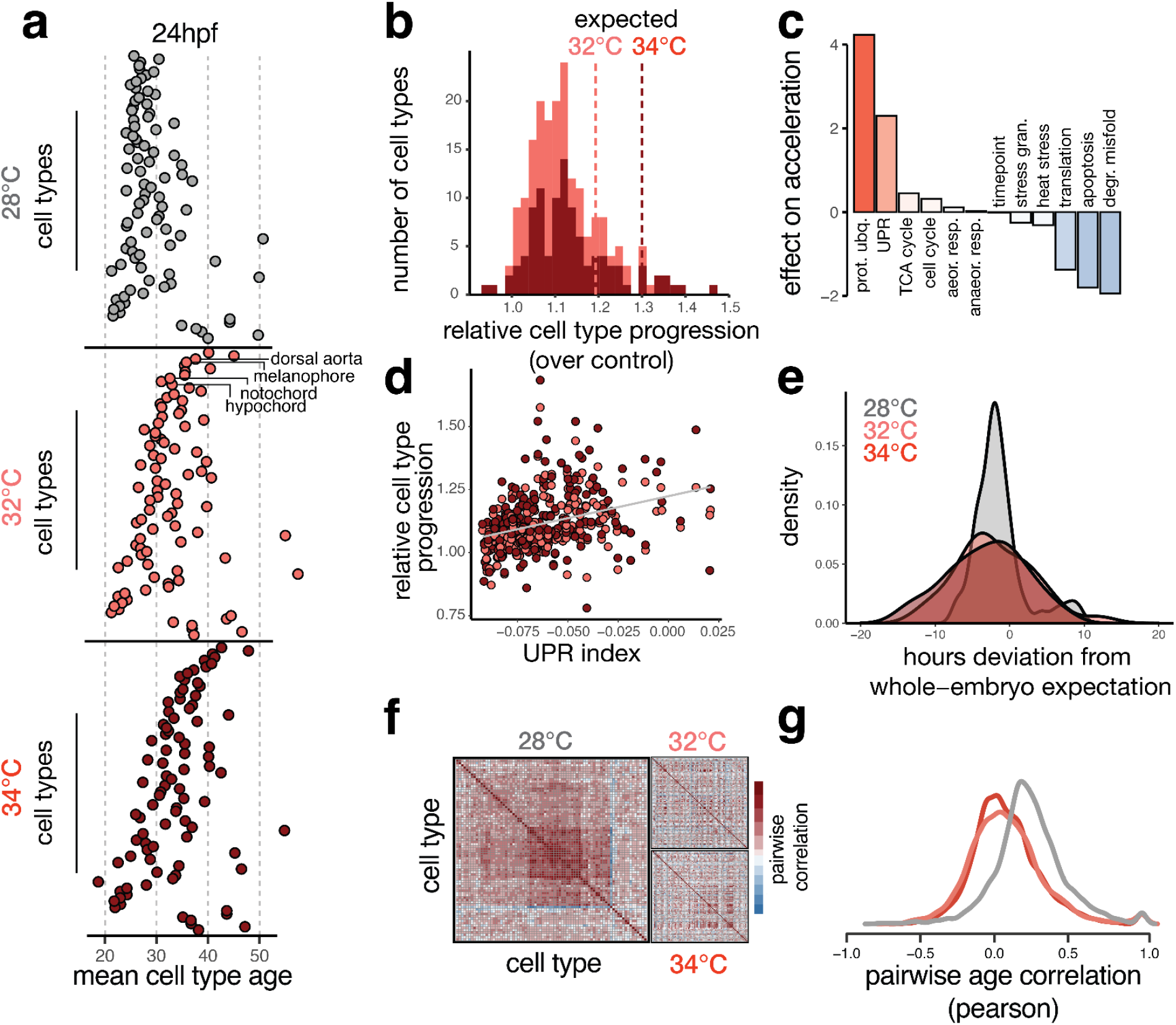
Temperature introduces asynchrony in developmental rate across cell types. **a**, Dotplots showing the transcriptional ages of all cell types at 24 hpf, faceted by temperature. Cell types on the y-axis are ordered by their relative acceleration at 34°C, highlighting the most sensitive cell types near the top, and insensitive types near the bottom. Cell type ordering is the same for all temperatures; specific examples are indicated with labels. **b**, Histogram showing distributions of relative cell type progression at each temperature, with vertical lines showing the relative progression expected for the whole embryo. **c**, Barplot showing the effect sizes for expression levels of several cellular processes related to metabolism, protein folding, proliferation, and stress response in an additive model predicting relative progression of the cell type at high temperature. **d**, Scatterplot showing basal levels of the unfolded protein response in each cell type against its relative progression, revealing a positive trend for both temperatures. **e**, Density histogram showing increased variance in developmental stage for embryos raised at elevated temperature. **f**, Heatmaps showing pairwise correlation coefficients of transcriptional age for all cell types in the embryo; at 28°C, most cell type pairs are positively correlated across individual embryos, whereas this correlation structure is diminished at elevated temperatures. **g**, Density histogram of pairwise correlation values at each temperature, summarizing the loss in correlation structure see in panel (F).

Given that some cell types were more developmentally accelerated than others, we next sought to directly quantify the synchrony of development across cell types. We first assessed variation in developmental stage at the embryo level by examining pseudostage deviations at each temperature; both temperatures increased variability in developmental stage (**Fig. 3e**). We also found that coordinated developmental timing between cell types could be observed in the covariance structure of cell type ages across all embryos (**Fig. 3f**). As expected, there was a positive correlation in age between all cell type pairs in control embryos (average r = 0.24) (**Fig. 3g**). However, we found that the covariance structure between cell types was disrupted in both of our elevated temperature conditions; the correlation of transcriptional age between cell type pairs was reduced by three-fold on average (**Fig. 3g**). Together, these results highlight how temperature introduces a non-uniform increase in developmental rate across different cell types and disrupts the developmental synchrony between cell types. Together, these results highlight an additional source of temperature-induced variability during development: asynchrony in developmental rate emerging from cell type-specific sensitivity to temperature.

Without an obvious anatomical or lineal relationship uniting highly accelerated cell types, we wondered whether any underlying molecular processes could explain differential developmental acceleration. We hypothesized that differences in baseline levels of 11 major cellular processes such as cell cycle, protein synthesis, metabolism, or stress response might be associated with the differences in relative developmental rate increases among cell types. To test this, we modeled the developmental rate increases across cell types as a function of baseline (during normal development at standard temperature) expression of genes important for each process (**Fig. 3c, inset**). Cell types expressing higher levels of genes involved in protein synthesis, apoptosis, and degradation of misfolded protein showed less temperature-dependent acceleration of development (**Fig. 3c**), slowing down relative to whole-embryo expectation. Cell types with high expression of genes involved in protein ubiquitination and the unfolded protein response (UPR) showed more temperature-dependent acceleration of development than those with lower expression of these processes. That temperature-dependent acceleration across all cell types could be partially explained by baseline levels of UPR suggests a role for ER stress in modulating developmental rate under normal conditions (**Fig. 3d**). Activation of UPR during normal development has been observed in both the notochord and the hatching gland ^27–29^ in fish, as well as during cell differentiation and growth in other models ^30,31^, but has not been previously linked to developmental rate.

### The unfolded protein response buffers notochord-specific temperature sensitivity

To explore whether transcriptional responses to temperature, similar to changes in cell abundances and developmental age, were non-uniform across cell types, we analyzed genes with differential expression at higher temperatures. We identified several modules of genes that tended to be upregulated in response to elevated temperature (**Extended Data Fig. 4a, Supplementary Table 4**). These modules were enriched for genes related to cell cycle, chromatin organization, transcription, ion transport, and protein processing in the ER, but not the canonical heat-shock response (**Extended Data Fig. 4a, Extended Data Fig. 1d, Supplementary Table 5**). In contrast to global activation of the heat-shock response ^23^, these temperature-responsive modules tended to be upregulated in a cell type-specific manner (**Extended Data Fig. 4b**). For example, only blood cells showed activation of the cell cycle module, while the notochord showed concomitant activation of translation and ER stress modules (**Extended Data Fig. 4b, boxed**). Of all the temperature-response modules, the ER stress gene cluster showed the broadest response across tissues.

The unexpected appearance of the UPR (**Fig. 3c**) and a gene module for translation and ER stress (**Extended Data Fig. 4a**) among cell type-specific responses to temperature led us to explore this phenomenon more deeply across cell types. We generated signature scores for translation and the UPR in all cell types in individual embryos and found that cells tended to express high levels of either one signature or the other, consistent with the UPR’s role in attenuating translation (**Fig. 4a, Supplementary Table 6**) ^32^. In contrast, we observe in the notochord a simultaneous upregulation of UPR and translation that is specific to elevated temperature conditions (**Fig. 4b, Extended Data Fig. 5a**). This exceptional upregulation of translation in the presence of UPR is pronounced to a lesser extent in the hypochord and pectoral fin bud; nearly all other cell types showed reductions in the number of cells expressing both signatures (**Fig. 4b**). Notably, the notochord showed temperature-dependent effects on cell abundance, as well as a marked acceleration of developmental age (**Fig. 2e, Fig. 3a, Extended Data Fig. 5b**). We wondered whether this exceptional upregulation of UPR and translation might be related to the notochord’s apparent sensitivity to temperature.

**Figure 4.**
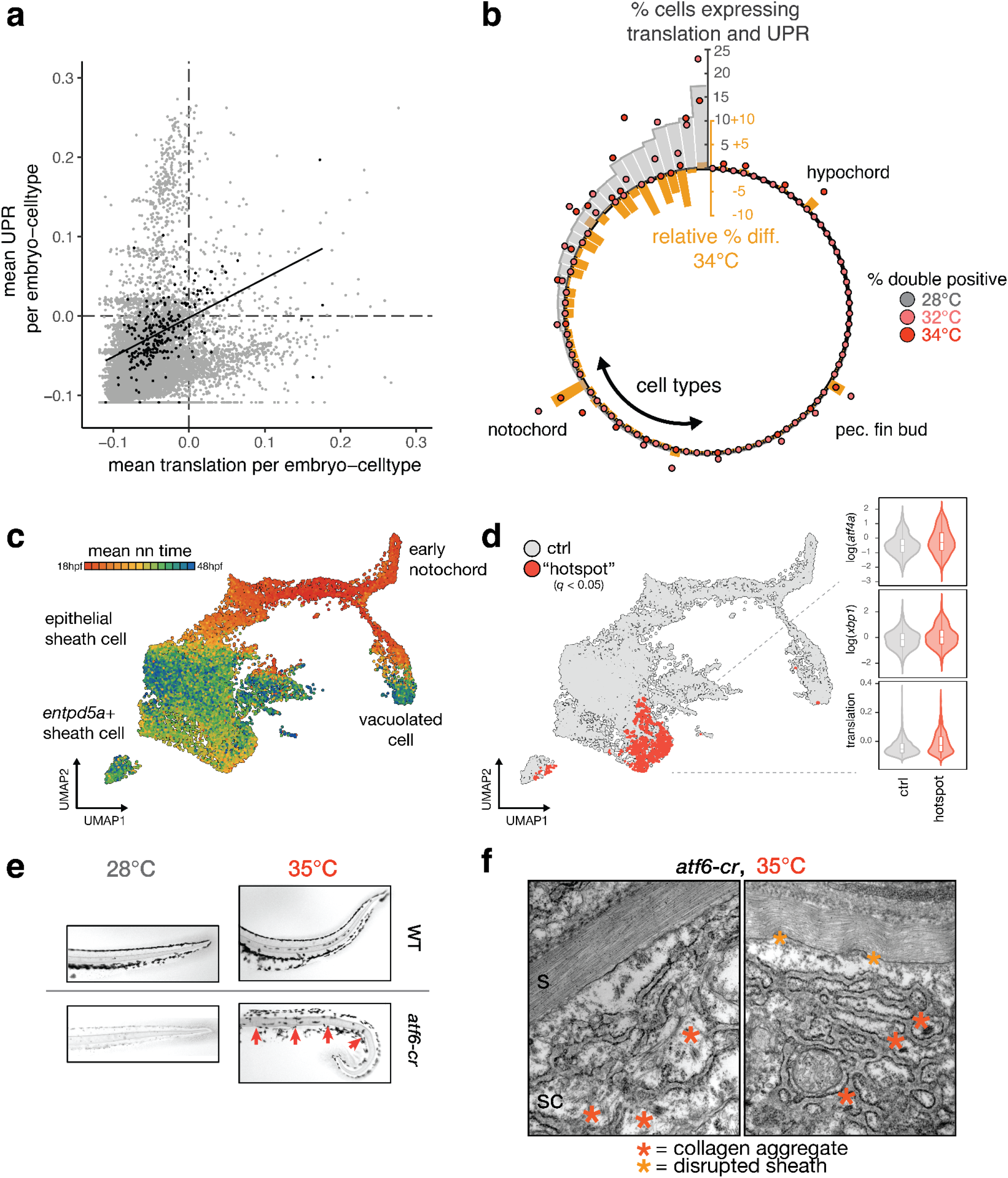
Exceptional regulation of the unfolded protein response underlies temperature sensitivity in the notochord. **a**, Scatterplot showing levels of translation signature and UPR signature across all embryo-cell types. Consistent with known translational attenuation by UPR, these processes are generally uncorrelated across cell types. The notochord (plotted as black dots) is an exception, with a positive correlation (Pearson’s r = 0.51) between these processes, and appreciable levels of both UPR and translation in a subset of embryos. **b**, Circular barplot showing occurrence of unusual translation + UPR, displayed as % double positive in each cell type (grey bars). Overlaid dots show raw % double positive cells for embryos raised at elevated temperature, colored pink and dark red for 32°C and 34°C, respectively. Orange bars show the relative change in the fraction of double positive cells in each cell type at elevated temperatures. For the large majority of cell types, this difference is negative. The notochord, hypochord, and pectoral fin bud are the sole exceptions, where double positive cells increase in response to temperature increase. **c**, UMAP showing co-embedded notochord cells from reference and temperature perturbation experiment, colored according to the mean time point label of nearest neighbors in the reference. Annotations for each cell type are indicated. **d**, UMAP showing sheath cells from temperature perturbation experiment with significant spatial bias using hotspot test (q < 0.05) in UMAP space (see Methods). Cells with a significant spatial base are shown in red. Inset shows levels of UPR markers and translation signature in these cells increasing with temperature. **e**, Images of Alcian-blue stained notochords (camera image in black-and-white) in the tails of wild-type and *atf6* crispant embryos raised at 28°C and 35°C. Examples of notochord defects (kinks, sheath collapse) in crispants raised at elevated temperature are indicated with red arrows (phenylthiourea was used for imaging 28°C crispants). **f**, Results of TEM of sheath cells in *atf6* crispants raised at elevated temperature, showing disrupted ER structure and aggregated collagen fibrils (red asterisks).

The notochord has unique mechanical and signaling functions, with specific cell sub-types carrying out these roles. These notochord sub-types are apparent in the reference dataset, including the vacuolated inner cells (*rab32a+, cav3+*), epithelial-like sheath cells (*col8a1+*), and pre-mineralization sheath cells (*entpd5a+*) (**Fig. 4c, Extended Data Fig. 5d**). To identify which of these types might be responsible for its temperature-sensitivity, we compared notochord cells from embryos raised in elevated temperatures with notochord cells from our wild-type reference dataset (**Fig. 4d**). We detected significant enrichment for cells from the elevated temperature condition in focal regions of the notochord UMAP, including a “hotspot” among the epithelial-like sheath cells, which expressed genes for translation and UPR at higher levels than other notochord subtypes (**Fig. 4e**). The primary function of the epithelial-like sheath cells is to produce, process, and package large amounts of collagen and ECM proteins that will ultimately be secreted to produce the extracellular notochord sheath, which is essential for the rigidity of the notochord and the proper formation of the embryonic body ^33^.

To test whether UPR buffers temperature-mediated ER stress in the developing notochord, we used CRISPR in F0 embryos to knock out the ER stress-sensing transcription factor Atf6 and examined notochord morphology using Alcian blue stain. In accordance with previous descriptions in medaka fish ^29^, we observed slight notochord defects at both 24 hpf and 48 hpf when fish were raised at standard temperature (**Fig. 4e**) and nearly all *atf6* crispants were viable past 24 hpf. We next tested if defects in *atf6* crispants were exacerbated when these embryos were raised at elevated temperature, especially given the proteostatic stress expected from temperature-dependent upregulation of translation in the notochord. Notochords from control fish raised at elevated temperature looked normal except for the usual bends, while *atf6* crispant fish showed kinks, bends, or deformations of the notochord sheath (**Fig. 4e**). These phenotypes evoke notochord mutants that affect the structural integrity of the sheath, such as *col8a1*, rather than the mutants affecting vacuolation, such as *rab32a* ^*34,35*^.

Because the sheath cells showed a potent upregulation of genes required for translation in response to increased temperature (**Fig. 4e**), we wondered whether this cell type was functioning properly in deposition of the notochord sheath. We examined the ultrastructure of sheath cells and the sheath using transmission electron microscopy. We confirmed that the notochord sheath showed increased disorder at 35°C in both wild-type and *atf6* crispant fish compared to controls raised at standard temperature (**Fig. 4f, Extended Data Fig. 5c**), suggesting that proper modification and organization of collagen is challenged at elevated temperature. Furthermore, we note a significant disruption of ER structure in this condition. The most striking difference in *atf6* mutants raised at high temperature was an accumulation of misfolded, non-secreted collagen fibers in the sheath cells themselves (**Fig. 4f**). Both protein aggregates and the severe disruption of ER structure in these fish suggest a failure of export, likely owing to the increased burden of protein synthesis at high temperatures combined with a lack of a properly functioning ER stress response. Taken together, these experiments show that in order for the notochord to structurally support the developing embryo, one of its cell types must operate near its proteostatic limits, requiring management by the UPR even at normal temperatures. Elevated temperature pushes the notochord sheath cells past their proteostatic limits, leading to protein aggregation, cellular defects, and a compromised notochord sheath that ultimately leads to a bent tail.

## Discussion

In characterizing cell type-specific responses to temperature in embryos, we identify previously overlooked sources of compromised developmental robustness. First, we find variability in the temperature-induced developmental acceleration across cell types, introducing asynchrony in the coordinated and timely traversal of developmental trajectories. This phenomenon is not unlike the consequence of genetic mutants causing stalling or mis-timed events of single cell types in the coordinated program of development ^36^. Unlike in these genetic mutants, elevated temperature does not induce stalling in any single cell type but rather introduces asynchrony across cell types; the degree of accelerations among cell types ranges from negligible to over 40% and correlated progression between cell types is lost. In sum, elevated temperature challenges the robustness of the developmental program by introducing non-uniform acceleration of developmental timing across cell types, breaking synchrony across the embryo.

Cell type-specific sensitivity to temperature is evident in the altered cell composition of embryos raised under stress. The notochord was especially sensitive, owing to the burden of protein synthesis in epithelial sheath cells which, under high temperature conditions, accumulate misfolded protein. The developing notochord depends on high levels of protein synthesis ^37^, ER chaperone function ^38^, and specialized secretion ^39^ to construct a well-ordered, collagen-rich sheath. This results in considerable stress on the ER and physiological activation of the UPR. The surprising upregulation of translation in these cells despite presumed translational attenuation by the UPR suggests that sheath cells might bypass translation regulation to achieve their exceptional protein synthesis capacity ^40^. Several other cell types with high demands of protein expression also depend on stress responses during normal development; the UPR is activated in the hatching gland, periderm, and vascular smooth muscle and via the heat shock response (HSR), cytoplasmic chaperones are activated in red blood cells, fast muscle, and cardiomyocytes. While the consequences of excessive burden on protein homeostasis in cells are evident in the aggregation associated with neurodegenerative diseases ^41^, we note the impact of this burden during embryogenesis, when even a transient stress may irreversibly perturb normal development. We propose that cell types with high demands on proteostasis during normal development may be more susceptible to further perturbations of homeostasis by temperature or other environmental stressors. Further exploration into the differential demands placed by each cell type on ancient and general molecular processes may shed new light on how stress on a molecular pathway in a specific cell type generates phenotypes at the tissue, organ, or organismal scale.

## Methods

### Animal rearing, staging and stocks

Staging followed Kimmel ^17^ and fish were maintained at ∼28.5°C under 14:10 light:dark cycles. Fish stocks used were: wild-type AB. Embryos were shifted to elevated temperatures at 6 hpf, and remained at these temperatures until sampling. All procedures involving live animals followed federal, state and local guidelines for humane treatment and protocols approved by the Institutional Animal Care and Use Committee (protocol #2997-01) of the University of Washington.

### Preparation of barcoded nuclei from individual embryos

Individual zebrafish embryos (24 to 36 hpf) were manually dechorionated and transferred to separate wells of a 96-well V-bottom plate containing 75 uL of 1X TrypLE + 2mg/mL Collagenase P (Millipore Sigma, cat. 11213865001). Embryos were then dissociated manually at 30°C by pipetting every 5 minutes for about 20 minutes until no visible chunks were present under a dissecting scope. Stop solution (dPBS, 5% FBS) was then added to each well and cells were spun down at 600xg for 5 min. Cells were then rinsed 1X in cold, 1X dPBS and spun down again. After rinses, supernatant was fully removed and cells were resuspended in 50 uL of CLB (Nuclei buffer, 0.1% IGEPAL, 1% SuperaseIn RNase Inhibitor (20 U/μL, Ambion), 1% BSA (20 mg/ml, NEB)) + hash oligos (1 uM, IDT) and incubated for 3 min on ice to liberate nuclei and integrate hash barcodes. To each well, 200 ul of ice cold, 5% Paraformaldehyde was added to each well. After an additional round of mixing, nuclei were fixed on ice for 15 minutes. All wells were then pooled together in a 15 mL conical tube and spun down for 15 min at 750xg. Supernatant was decanted and cells rinsed in 2mL of cold NBB (Nuclei Buffer, 1% BSA, 1% SuperaseIn) at 750xg for 6 min. Supernatant was then removed and cells were resuspended in 1mL of NBB and flash frozen in LN2 and stored at -80.

### sci-RNA-seq3 library construction

The fixed nuclei were processed similarly to the published sci-RNA-seq3 protocol ^22^. For paraformaldehyde fixed cells, frozen fixed cells were thawed on ice, spun down at 750xg for 6 min, and incubated with 500ul NBB (Nuclei buffer, 1% BSA, 1% SuperaseIn) including 0.2% Triton X-100 for 3 min on ice. Cells were pelleted and resuspended in 400ul NBB. The cell suspension was sonicated on low speed for 12s. Cells were then pelleted at 750xg for 5 min prior to resuspension in NB + dNTPs. The subsequent steps were similar with the sci-RNA-seq3 protocol (with paraformaldehyde fixed nuclei) with slight modifications: (1) We distributed 25,000 fixed cells (instead of 80,000 nuclei) per well for reverse transcription. (2) Centrifugation speeds for all spins were increased to 750xg.

### Sequencing, read processing and cell filtering

Libraries were sequenced on an Illumina Novaseq 6000 (S4 200 cycle kit) with sequencing chemistries compatible with library construction and kit specifications. Standard chemistry: I1 - 10bp, I2 - 10bp, R1 - 28bp, R2 - remaining cycles (> 45). Read alignment and gene count matrix generation was performed using the Brotman Baty Institute (BBI) pipeline for sci-RNA-seq3 (https://github.com/bbi-lab/bbi-sci). After the single cell gene count matrix was generated, cells with fewer than 200 UMIs were filtered out. For mitochondrial signatures, we aggregated all reads from the mitochondrial chromosome. Each cell was assigned to a specific zebrafish embryo based on the total hash count (hash_umi >= 5) and the enrichment of a particular hash oligo (hash enrichment cutoff >= 3).

### Count matrix pre-processing

Transcript by cell count matrix was pre-processed using a standard Monocle3 workflow:

*estimate_size_factors()* -> *detect_genes(min_expr = 0*.*1)* -> *preprocess_cds()* with 100 principal components for whole-embryo and 50 principle components for subsets, *align_cds(residual_model_formula_str = “∼log10(n*.*umi)”), reduce_dimension(max_components = 3, preprocess_method = ‘Aligned’)*, and finally *cluster_cells(resolution = 1e-4)*.

### Cell annotation by projection of query data to reference

Cell type annotations were assigned via a label transfer procedure from the reference dataset (**see accompanying manuscript, Saunders et al. 2022**) to the query temperature-treated dataset. Briefly, the PCA rotation matrix, log10(n.umi) batch correction linear model, and non-linear UMAP transformation from the reference data were computed and saved for subsequent transformation of query data. The query temperature-treated dataset was then projected into the reference space using the following procedure: coefficients to transformed gene expression values into PCA loadings were applied, the linear batch correction was then applied to remove effects of total UMIs per cell, and, lastly, the UMAP transformation was applied to the batch-adjusted PCA loadings to project query data into the reference coordinate space. A similar overall approach is used in Andreatta *et al*.^42^

After an initial transfer of the ‘major group’ label (four possibilities: mesoderm, mesenchyme-fin, periderm + other, CNS) in the global space of all cells, cells that gained major group labels were projected in a sub-space corresponding to that major group. In this sub-space, finer resolution annotations (germ layer, tissue, broad cell type, sub cell type) were transferred using the majority vote of reference neighbors (k=10).

### Estimation of transcriptional age for each cell and cell type

Alongside the process for transferring cell type annotation labels as above, each cell from the query temperature-treated dataset was assigned an estimated timepoint label (mean_nn_time) by taking the average sample timepoint of its 10 nearest neighbors in the developmental reference dataset. For each embryo, we bulked and averaged all cells of the same type to compute a cell type-specific “transcriptional age.”

### Comparison of embryo pseudostage with conventional staging

A linear model for the effects of temperature on developmental stage from Kimel et al was used to generate stage expectations: h = H * (0.055T - 0.57), where h = hours of development to reach the stage at 28.5°C and H = hours of development at temperature T.^17^

### Additive model for cell type-specific acceleration

Signatures scores were computed using the *aggregate_gene_expression()* function in Monocle3, using the log-normalized average counts across gene sets representing several biological processes (**Supplemental Table 6**). To assess the baseline contributions of each process to acceleration by temperature, signature scores were computed for embryos raised at the standard 28°C. A linear model was used (vglm from VGAM package ^43^) to examine the relationship of each signature with relative developmental acceleration, and coefficients for each process were compared.

### CRISPR/Cas9 mutagenesis

gRNAs were designed using either the IDT or CRISPOR ^44^ online tool. gRNAs were synthesized by IDT as crRNAs, annealed to tracrRNAs and assembled into RNPs with spCas9 protein (IDT). Preparation and subsequent injection of RNPs was performed as previously described;^45^ two gRNAs were simultaneously injected for *atf6*.

### Transmission Electron Microscopy

Fish were euthanized then fixed in sodium cacodylate buffered 4% glutaraldehyde overnight at 4°C. Embryos were stained in 2% osmium tetroxide for 30 min, washed, and then stained in 1% uranyl acetate overnight at 4°C. Samples were dehydrated with a graded ethanol series then infiltrated with a 1:1 propylene oxide:Durcupan resin for 2 hours followed by fresh Durcupan resin overnight and flat embedded prior to polymerization. Blocks of posterior tissue, from yolk extension to tail, were thin sectioned on a Leica EM UC7 and sections imaged on a JEOL 1230 transmission electron microscope.

### Notochord staining

Alcian blue staining followed an online procedure (SDB Online Short Course, Zebrafish Alcian Blue), except that embryos were raised in 1-phenyl-2-thiourea to suppress pigment formation instead of bleaching.

### Differential gene expression analysis

To capture biological variation in gene expression between stage-matched embryos, cells of each type were first pseudobulked (sums of size-factor-normalized gene counts) by embryo. Using this embryo-level pseudobulk data as input, differentially expressed genes were determined in each cell type using the fit_models() function in Monocle3. The model formula included additive terms for temperature (as a factor), a spline of embryo pseudostage (degrees of freedom = 3), and the total number of cells. Global analysis of temperature-dependent gene expression was performed by extracting temperature coefficients from the model output, filtering for genes with significant q-values (q < 0.05). Temperature coefficients in each cell type were combined into a single matrix, and processed analogously to the find_gene_modules() function in Monocle3 (ref), which uses a transposition of the typical gene x cell matrix to group genes with similar patterns across cells, rather than grouping cells. Temperature coefficients (no normalization) were pre-processed with PCA and dimensionality reduction was performed with UMAP (n_neighbors = 25L, min_dist = 0.2). Clusters of genes with similar temperature responses across cell types were identified using the cluster_cells() function.

### Cell abundance and variance statistics

Changes in the proportions of each cell type were assessed by first counting the number of each annotated cell type in each embryo. To control for technical differences in cell recovery across embryos, “size factors” by dividing the total number of cells recovered from an embryo by the geometric mean of total cell counts across all embryos. The number of cells of each type recovered from each embryo were then divided by that embryo’s size factor.

Normalized counts for each cell type *i* at time *t* were then compared across genotypes using a generalized linear model defined by the equations:

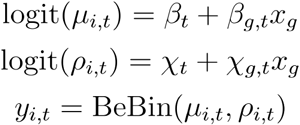

Where *y*_*i,t*_, the normalized counts of cell type *i* at time *t* is modeled as a beta-binomially distributed random variable with mean *μ*_*i,t*_ and “litter effect” *ρ*_*i,t*_ (i.e. overdispersion with respect to the binomial distribution). We modeled both parameters of the beta binomial response as a function of genotype, reasoning that crispants might exhibit greater variability than wild-type embryos. The binary indicator variable *x*_*g*_ denotes whether gene *g* is knocked out in each embryo. Separate models for each temperature in each cell type and at each pseudostage bin were fit using the VGAM package ^43^. Significance of knockout effects in each model were assessed by Wald test on *β*_*g,t*_.

### Hotspot analysis

The local spatial statistic Getis-Ord index (G_i_) was used to identify statistically significant regions of the UMAP embedding that were enriched or depleted of perturbed cells. A high-value G_i_ indicates that a perturbed cell is surrounded by other cells with the same perturbation, whereas a G_i_ close to zero indicates that a perturbed cell is surrounded by cells with other perturbation labels. A G_i_ was calculated for each cell’s local neighborhood (k=15) using the “localG()” function in the spdep package. This returns a z-score that indicates whether the observed spatial clustering is more pronounced than expected by random. Multiple testing correction was done using a Bonferroni correction. Areas of the UMAP where a given perturbation is enriched are called “hot spots.”

## Acknowledgements

We thank Nola Klemfuss and the Brotman Baty Institute Advanced Technology Lab for support with sequencing and data processing. We also thank Frank Steemers and Fan Zhang for additional sequencing support. We thank Ed Parker for assistance with TEM.

## Author contributions

M.W.D., L.M.S., and C.T. designed the study. M.W.D., L.M.S., and S.S. performed sci-RNA-seq3 experiments.

M.W.D. performed computational analyses with M.D, B.E. and C.T. D.K. performed notochord staining.

M.W.D. and C.T. wrote the manuscript with input from all coauthors. J.S., C.Q., D.R., and D.K. contributed methods, supervision, and edited the manuscript. C.T. supervised the project.

## Data Availability

The datasets generated and analyzed during the current study are available in the NCBI Gene Expression Omnibus (GEO) repository: GSE202294.

## Code Availability

Pipelines for generating count matrices from sci-RNA-seq3 sequencing data are available at https://github.com/bbi-lab/bbi-dmux and https://github.com/bbi-lab/bbi-sci. Analyses of the single cell transcriptome data were performed using Monocle3, which was updated to include methods from this study; a general tutorial can be found at http://cole-trapnell-lab.github.io/monocle-release/monocle3. Additional scripts for performing data processing, statistical analysis and generating plots will be made available via a Github repository (https://github.com/cole-trapnell-lab/sdg-zfish).

## Figures

**Extended Data Fig. 1.**
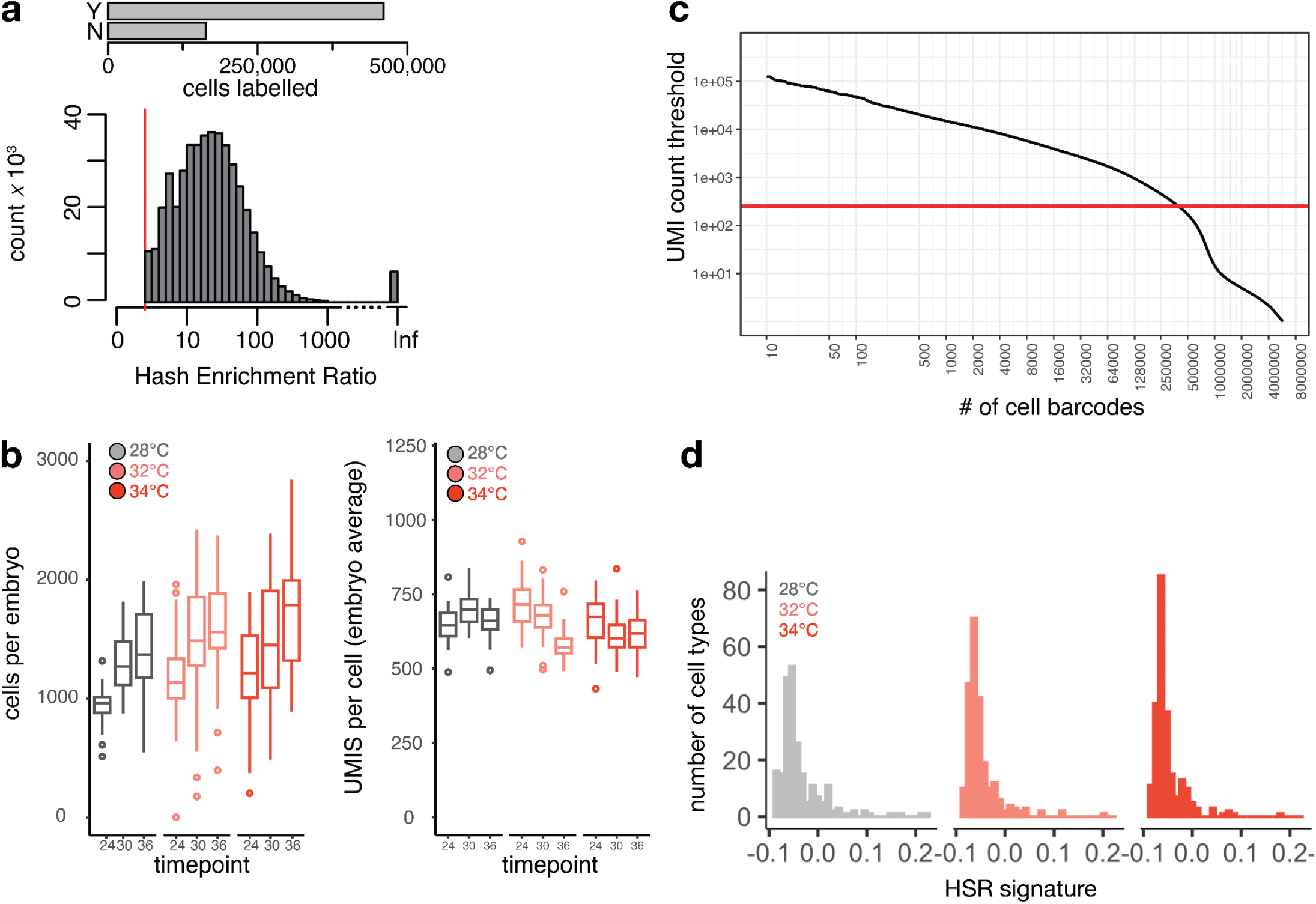
Individual-level capture of high-quality single cell transcriptomes. **a**, Barplot showing the number of individual cells that pass (Y = yes, N = no) a Hash Enrichment Ratio (count of most abundant hash relative to second-most abundant) threshold of 3. Below is a histogram showing the full distribution of Hash Enrichment Ratios; cells with only a single hash captured are labeled as ‘Inf.’ **b**, Boxplots showing the number of cells in each embryo (left) and number of UMIs per cell in each embryo (right), faceted by sample time point and colored by temperature treatment. **c**, Knee plot showing the threshold for UMI count (red line) in called cells. **d**, Histogram showing no systemic activation of the heat shock response (HSR) across cell types at any of the temperatures profiled. While the total number of cell types with negligible HSR levels increases with temperature, nearly all cell types should have HSR signature values greater than 0 in the event of global activation of the HSR.

**Extended Data Fig. 2.**
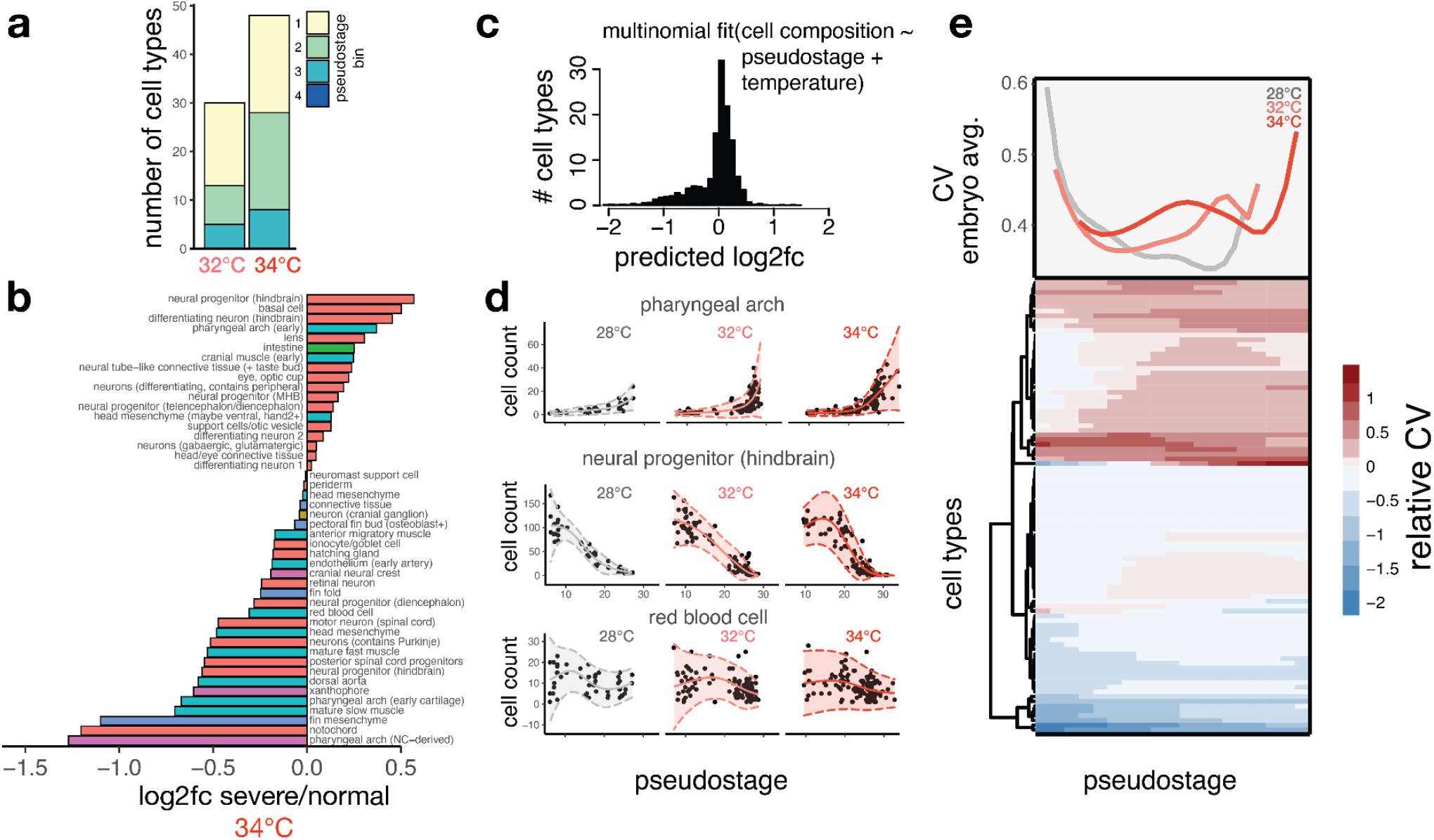
Temperature increases variability in specific cell types and stages of development. **a**, Stacked barchart showing how the number of cell types with significant abundance changes increases with temperature; this phenomenon held for all pseudostage bins (in color) except bin 4, which lacked sufficient control embryos to analyze statistically. **b**, Barplot showing the log2 fold change in abundance for all cell types in 34°C embryos with severe phenotypes relative to those that were phenotypically normal. **c**, Histogram showing changes in cell composition resulting from 34°C treatment, as predicted by multinomial model. **d**, Scatterplots showing raw data (black points) and model outputs of mean (solid line) and variability (ribbons with dotted line) for three temperature-sensitive cell types. **e**, Heatmap of each cell type showing relative increases in observed variability compared to variability expected during wildtype development through developmental time (pseudostage bins); shades of red indicate increased variability while white to blue shades show neutral or decreased variability. The upper panel shows the average CVs for each temperature across all stages, highlighting pseudostages with increased variability at higher temperatures.

**Extended Data Fig. 3.**
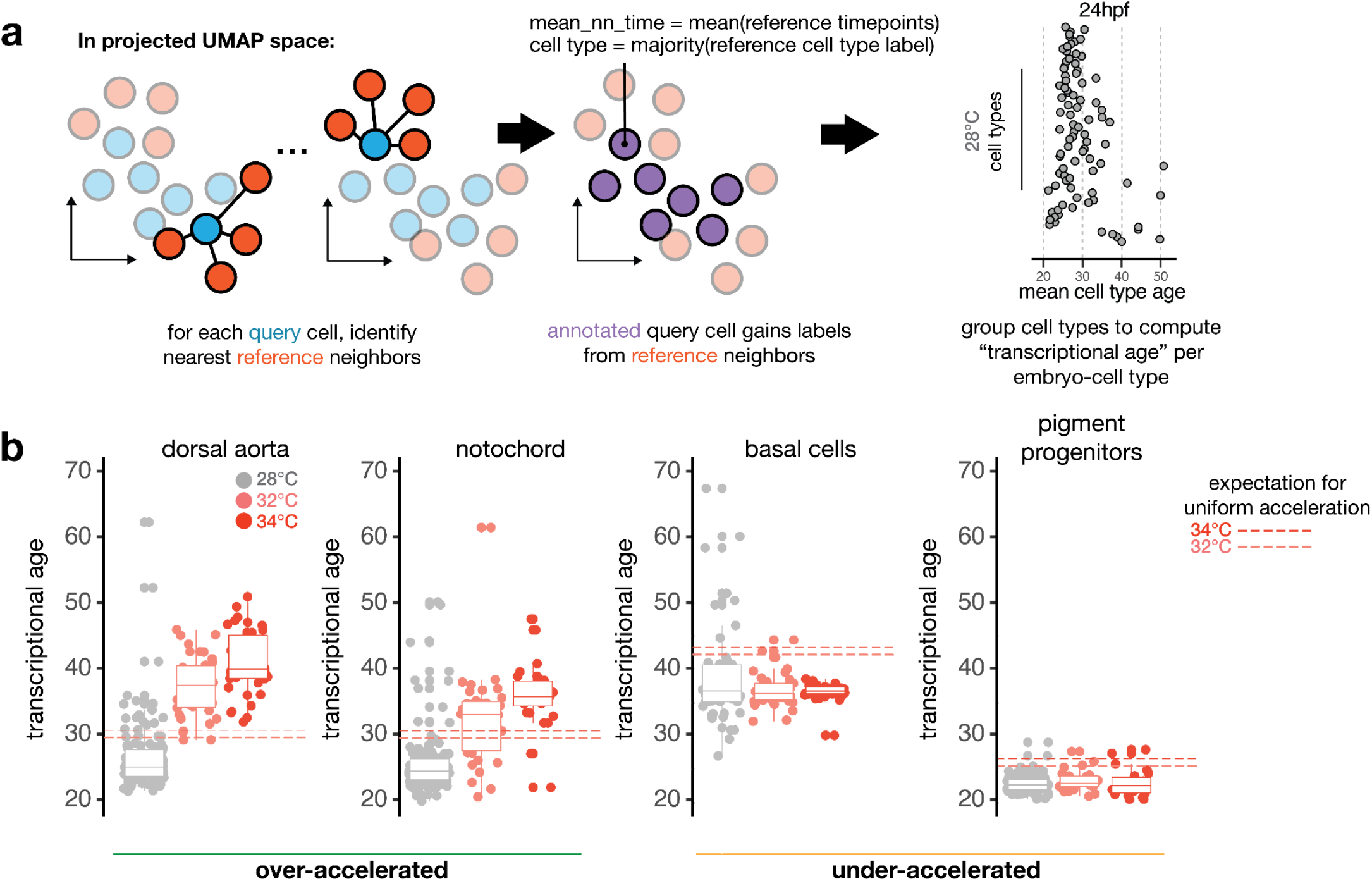
Temperature accelerates development non-uniformly across cell types. **a**, Schematic showing method for computing transcriptional age of each cell through nearest-neighbor averaging of time point labels in the reference. **b**, Boxplots showing individual examples of cell types with greater-than-expected acceleration (left) and less-than-expected acceleration (right), with whole embryo expectation from Kimmel *et al* for each temperature shown as dotted lines.17

**Extended Data Fig. 4.**
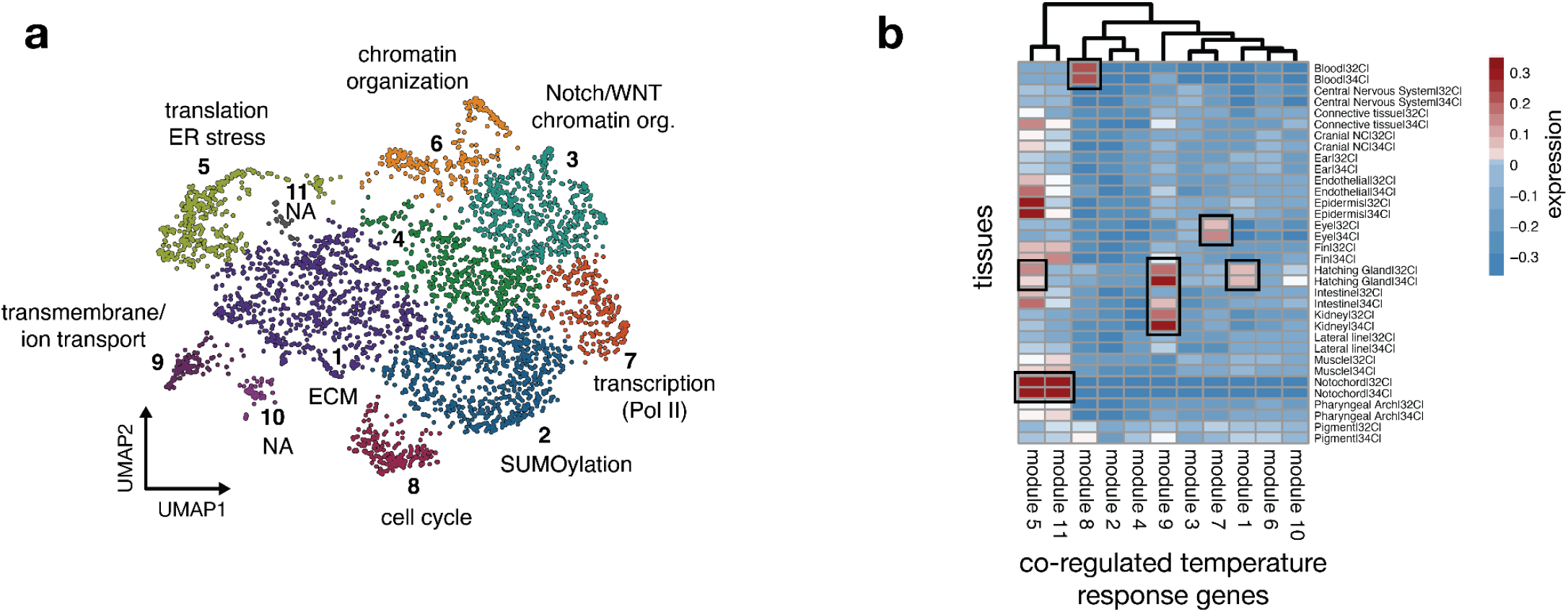
Transcriptional responses to elevated temperature tend to be cell type specific. **a**, UMAP of all genes with significant temperature-dependent transcriptional changes (see Methods). Each point represents an individual gene, and genes with similar responses (coefficient in differential expression model after accounting for stage) across cell types are grouped together. Each cluster is labeled with a different color, and a representative GO term for the group of genes in each cluster is indicated. Clusters in which gene groups returned no significant GO terms are labeled ‘NA’. **b**, Heatmap of expression levels for each gene module in (a) as an average for each cell type at elevated temperatures. Consistent with the heterogeneous grouping of temperature responses in (a), elevated module expression tends to be cell type-specific (examples boxed). Module 5, which contains genes related to translation and ER processing, has the most broad response across cell types, but the magnitude of the response is greatest within the notochord.

**Extended Data Fig. 5.**
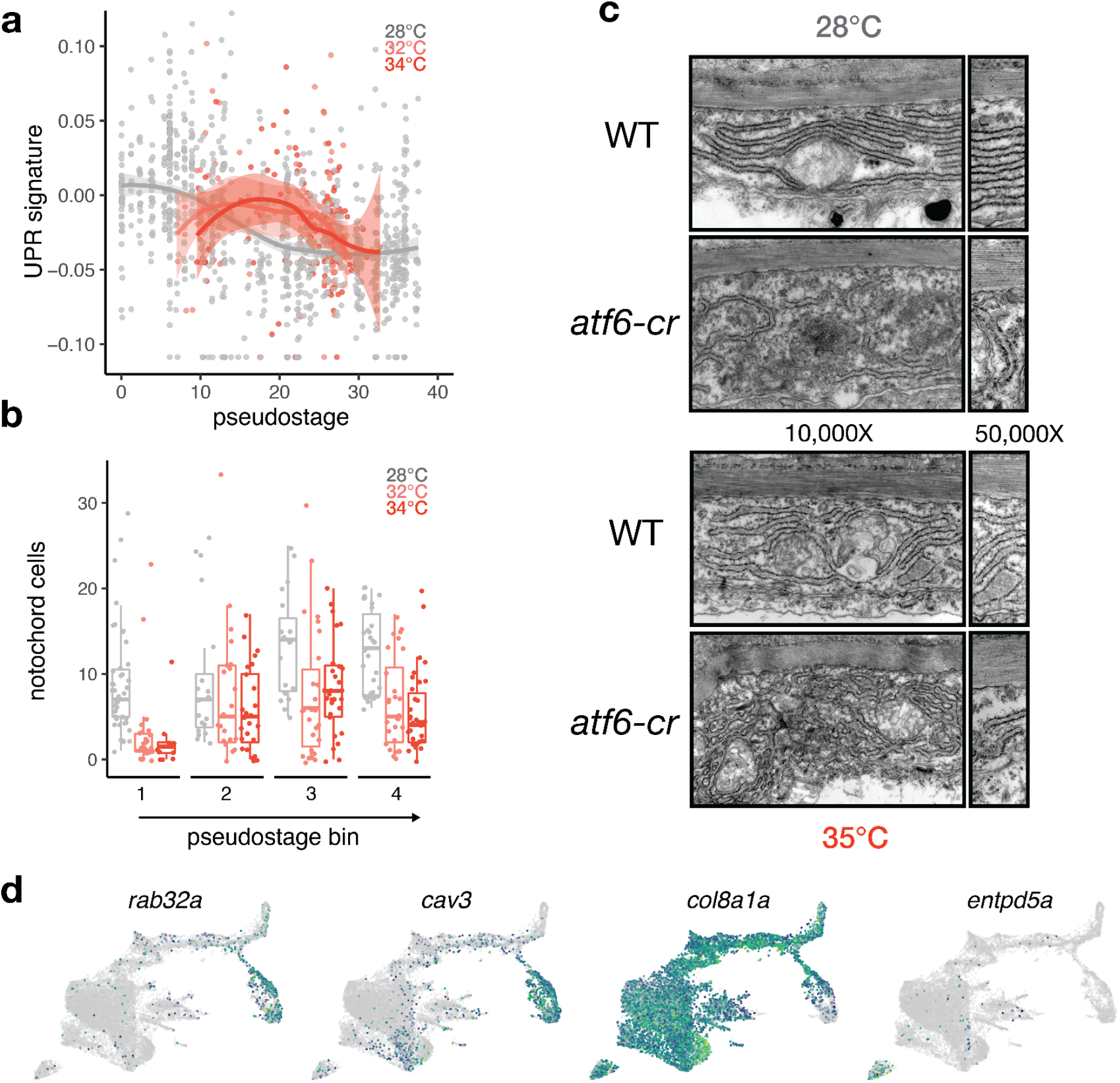
Burdens on proteostasis in sheath cells sensitize the notochord to stress. **a**, Boxplot showing marked reduction of notochord cells in embryos raised at high temperature, with raw counts in control and temperature-treated embryos shown as points. **b**, Scatterplot showing levels of UPR signature in the notochord cells of all individual embryos, colored by temperature. Embryos raised at higher temperature show higher levels of UPR signature at nearly all profiled stages. Lines and ribbons are derived from loess fits across pseudostage. **c**, TEM of sheath cells in wild-type and *atf6* crispants, showing effects in both untreated and temperature-treated embryos. ER structure is perturbed in *atf6* crispants, and severe sheath defects are visible in the temperature-treated *atf6* crispants. **d**, Marker gene plots showing expression of genes defining notochord sub-types (*rab32a* and *cav3* in vacuolated cells; *col8a1a* in sheath cells; *entpd5a* in pre-mineralization sheath cells).

